# Evolutionary differences in the ACE2 reveals the molecular origins of COVID-19 susceptibility

**DOI:** 10.1101/2021.03.25.437113

**Authors:** Ryan R. Cheng, Esteban Dodero-Rojas, Michele Di Pierro, José N. Onuchic

## Abstract

We explore the energetic frustration patterns associated with the binding between the SARS-CoV-2 spike protein and the ACE2 receptor protein in a broad selection of animals. Using energy landscape theory and the concept of energy frustration—theoretical tools originally developed to study protein folding—we are able to identify interactions among residues of the spike protein and ACE2 that result in COVID-19 resistance. This allows us to identify whether or not a particular animal is susceptible to COVID-19 from the protein sequence of ACE2 alone. Our analysis predicts a number of experimental observations regarding COVID-19 susceptibility, demonstrating that this feature can be explained, at least partially, on the basis of theoretical means.

## Introduction

The coronavirus disease 2019 (COVID-19) caused by the severe acute respiratory syndrome coronavirus 2 (SARS-CoV-2) has affected the lives of millions of people in a worldwide pandemic. The hallmark of COVID-19 is its high degree of contagiousness between individuals.

SARS-CoV-2 is believed to gain entry in to the host cell through its interaction with the Angiotensin-converting enzyme 2 (ACE2) receptor on the host cell surface(Zhou et al., 2020), similar to SARS-CoV-1(Li et al., 2003). Recently, the structure of the SARS-CoV-2 viral spike glycoprotein bound to the human ACE2 receptor was determined using X-ray crystallography (Wang et al., 2020), providing a crucial starting point for any molecular modeling of the viral interaction with ACE2. The structure of this crucial complex has also been independently determined using cryo-EM(Yan et al., 2020) and X-ray crystallography(Shang et al., 2020).

What are the molecular interactions that give rise to interaction specificity between the viral spike and ACE2 receptor? Hints at the molecular origins of COVID-19 susceptibility can be found analyzing the susceptibility of different organisms to the coronavirus. The ACE2 receptors are found in a diverse span of the animal kingdom, including mammals, birds, and aquatic life, which have varying degrees of COVID-19 susceptibility(BS et al., 2021; Gautam, Kaphle, Shrestha, & Phuyal, 2020; Goldstein, 2020; Kumakamba et al., 2020; Muñoz-Fontela et al., 2020; Mykytyn et al., 2020; Oude Munnink et al., 2021; Palmer et al., 2021; Shi et al., 2020; Sia et al., 2020; Sit et al., 2020).

For example, it is known that mice are immune to COVID-19 while on the contrary the Bronx Zoo tiger Nadia(Goldstein, 2020) had tested positive for COVID-19 and also exhibited many of the symptoms observed in infected humans. To date, a number of animals have been classified as either being susceptible or immune to COVID-19(BS et al., 2021; Gautam et al., 2020; Goldstein, 2020; Kumakamba et al., 2020; Muñoz-Fontela et al., 2020; Mykytyn et al., 2020; Oude Munnink et al., 2021; Palmer et al., 2021; Shi et al., 2020; Sia et al., 2020; Sit et al., 2020).

The evolutionary divergence in the sequences of the ACE2 receptor found in different organisms can be related to such susceptibility of infection(Becker et al., 2020; Damas et al., 2020; Frank, Enard, & Boyd, 2020; Lam et al., 2020; Luan, Lu, Jin, & Zhang, 2020; Martínez-Hernández et al., 2020; Melin, Janiak, Marrone, Arora, & Higham, 2020). Here, we explore the molecular mechanisms by which some sequence variants of the ACE2 receptor appear to confer resistance to infection by virtue of their reduced binding affinity to the viral spike protein. We examine the sequences for ACE2 receptor across a selection of 63 representative animals and identify the residue interactions that are responsible for the reduced binding affinity, and thus COVID-19 resistance, using the concept of energetic frustration from the theory of protein folding (Onuchic, Luthey-Schulten, & Wolynes, 1997; Onuchic & Wolynes, 2004). Here, energetic frustration refers to unfavorable interactions between residues in a given protein structure that cannot be mitigated without structural rearrangement or residue level mutations. In the context of the ACE2/spike complex, frustrated interactions between residues of the ACE2 and the spike glycoprotein can also exist for a given structure of the protein complex.

The rarity of kinetics traps observed in the folding of proteins indicates that, in general, proteins do not exhibit a high amount of energetic frustration, which would instead create those kinetic traps(Onuchic et al., 1997; Onuchic & Wolynes, 2004). While folding kinetics suggest proteins to be “minimally frustrated”, some local frustration may be present; for example, local frustration could be functionally useful for tuning conformational dynamics. In protein complexes, a site frustrated in the monomeric protein may become less frustrated when the protein is bound to its counterparts, thus guiding specific association(Ferreiro, Hegler, Komives, & Wolynes, 2007; Parra et al., 2016).

In the case of the complex formed between the SARS-CoV-2 viral spike glycoprotein and the ACE2 receptor, we use changes in energy frustration as a proxy for changes in binding affinity. We use the crystal structure of the viral spike bound to the human ACE2 as a template to construct molecular models of the interaction between the viral spike and the ACE2 receptors of these different animals. We then calculate the changes in frustration with respect to the reference point constituted by the human ACE2 sequence (Ferreiro et al., 2007; Parra et al., 2016). This allows us to identify key residues of the ACE2 protein that appear to inhibit the binding of the spike glycoprotein and to predict whether or not a particular animal will be susceptible to COVID-19. The novelty of our approach and the key to our results resides in the fact that, while our procedure is based on the only input of the protein sequences of ACE2 receptor, our approach does incorporate a great deal of structural information about the protein complex, which is extracted from the crystal structure(Wang et al., 2020), and physico-chemical details about the energetics of protein folding and docking, which is synthetized in the energy function and results from decades of developments(Onuchic & Wolynes, 2004).

## Materials & Methods

### ACE2 protein sequences

The majority of ACE2 protein sequences were previously annotated from the genome assembles and sequencing data from the DNA Zoo Consortium(Dudchenko et al., 2017) (Available for download: https://www.dnazoo.org/post/the-first-million-genes-are-the-hardest-to-make-r). The full length protein sequences of the ACE2 proteins for mouse (*Mus musculus*), ferret (*Mustela putorius furo*), chicken (*Gallus gallus domesticus*), pig (*Sus*), duck (*Anas platyrhynchos*), Syrian golden hamster (*Mesocricetus auratus*), and mink (*Neovison vison*) were obtained from the Uniprot database(The UniProt, 2021) to supplement the sequences derived from the DNA Zoo. In total, 63 representative ACE2 sequences were used in our study (Table S1). For comparative analysis, a multiple sequence alignment was generated for the ACE2 sequences using Clustal Omega(Madeira et al., 2019).

### Homology Modeling

The crystal structure of the SARS-Cov-2 glycoprotein spike bound to the human ACE2 protein served as our starting template for constructing models of the glycoprotein spike bound to the ACE2 protein of other animals. We used the SWISS-MODEL (Waterhouse et al., 2018) to create homology models of 63 representative animals (Table S1) for a full list.

### Frustration Analysis

We performed an energy landscape analysis on the predicted ACE2-spike complex for different animals using the configurational frustration index(Ferreiro et al., 2007; Parra et al., 2016):

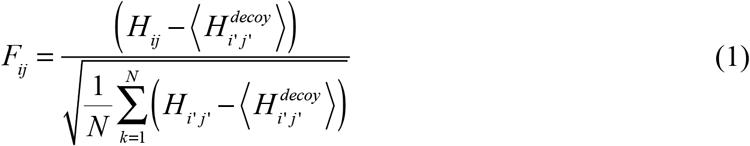

Here, *H*_ij_ represents the pairwise interaction energy between residues *i* and *j* in a given structure using the Associative Memory, Water Mediated, Structure and Energy Model (AWSEM)(Davtyan et al., 2012), a coarse-grained model widely used to study problems of protein folding and protein-protein association and assembly. The native energies *H*_ij_ are compared directly to *N* number of different configurational realizations between residues *i* and *j*, thereby generating a distribution of decoy energies with a mean of 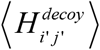 and a standard deviation of 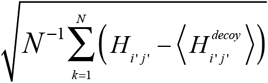. Hence, *F_ij_* is a type of Z-score that measures how favorable a particular pair of interactions are within a protein or protein complex with respect to a distribution of decoys. Frustrated (unfavorable interactions) are denoted by *F_ij_* < 0 while *F_ij_* > 0 are considered favorable; in particular, *F_ij_* < −1 is considered highly frustrated while *F_ij_* > 1 is considered minimally frustrated.

In our analysis, we found that it was useful to compare the configurational frustration between an interprotein residue pair with the same pair from the human ACE2-spike complex:

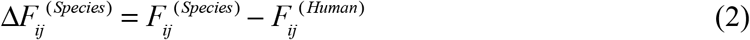

We find that 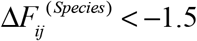 robustly identifies highly frustrated interactions that result in COVID-19 resistence. On the other hand, if all of the inter-protein residue interactions between the ACE2 receptor and the spike do not exhibit high levels of frustration (i.e., 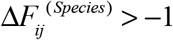 we identify that species as being highly susceptible to COVID-19. For completeness, if the most frustrated interprotein interactions fall between 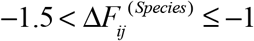 that species is predicted to be moderately susceptible.

### Evolutionary distance between ACE2 proteins

The Jukes-Cantor distance is used to quantify the evolutionary distance between aligned ACE2 proteins in our study: 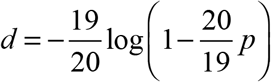, where *p* is the p-distance—i.e., the number of residue sites between two compared sequences that are different divided by the sequence length of the multiple sequence alignment.

## Results & Discussion

### Comparative frustration analysis of ACE2-spike complex for different species

By examining the comparative differences between the inter-protein interactions with respect to the human ACE2-spike complex, we are able to identify residue-interactions outliers that represent a significant disruption to the ACE2/spike interaction relative to the human ACE2-spike interaction.

Shown in Figure 1 are plots of our frustration analysis for mouse (*Mus musculus*) and tiger (*Panthera tigris*), which are used as representative examples of animals have been experimentally observed to be resistant(Muñoz-Fontela et al., 2020) and susceptible(Goldstein, 2020) to COVID-19, respectively.

**Figure 1.**
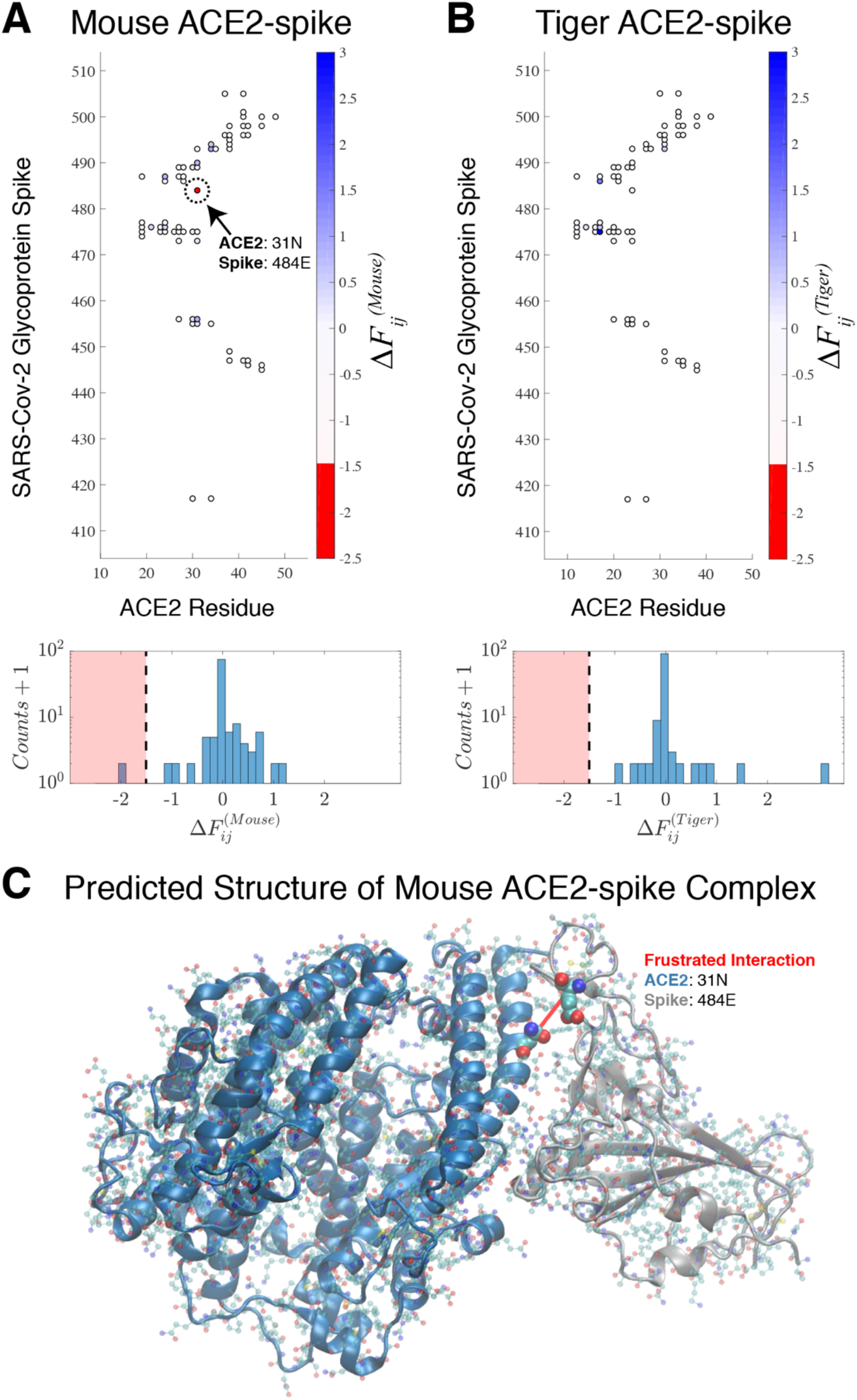
Comparative analysis of the frustration indices with respect to those observed in human ACE2-spike complex reveals outliers that indicate COVID-19 resistance. The configurational frustration index relative to the frustration in the human ACE2-spike complex is shown for (A) mouse (*Mus musculus*) and (B) tiger (*Panthera tigris*) on a contact map illustrating select contacts between the SARS-Cov-2 spike and the ACE2 protein. Corresponding histograms of the frustration index between all contacts between the ACE2 and spike protein are also shown. (A) and (B) are representative examples of animals that are resistant to COVID-19 and susceptible to COVID-19, respectively. Animals that resist COVID-19 appear to have frustrated outliers that represent highly unfavorable residue interactions compared to the human ACE2-spike complex (i.e., 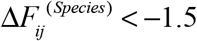). For the mouse, a single frustrated interaction between residue 31N of the ACE2 protein and 484E of the spike glycoprotein appears to confer COVID-19 resistance. (C) The frustrated interaction is plotted on the modeled 3D structure of the spike glycoprotein bound to the mouse ACE2 receptor.

We observe a single frustrated residue pair between 31N of the mouse ACE2 and 484E of the spike protein on the map of configurational frustration (Figure 2A/2C), which appears as an outlier in the histogram of 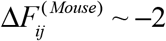. Similar highly frustrated outliers can be found for the other animals that are known to resist COVID-19 (Figure S1), such as chicken (*Gallus gallus domesticus*)(Shi et al., 2020) and duck (*Anas*)(Shi et al., 2020). We find that a threshold value of 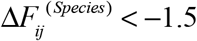 robustly identifies residue pairs that appear to confer COVID-19 resistence. Likewise, a histogram of 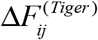 exhibits comparable levels of frustration to that of the human ACE2 and spike (Figure 1B)—similar findings are obtained for other animals with known susceptibilities to COVID-19 (Figure S2), such as white-tailed deer (*Odocoileus virginianus*)(Palmer et al., 2021), European rabbit (*Oryctolagus cuniculus*)(Mykytyn et al., 2020), and pig (*Sus scrofa*)(BS et al., 2021).

**Figure 2.**
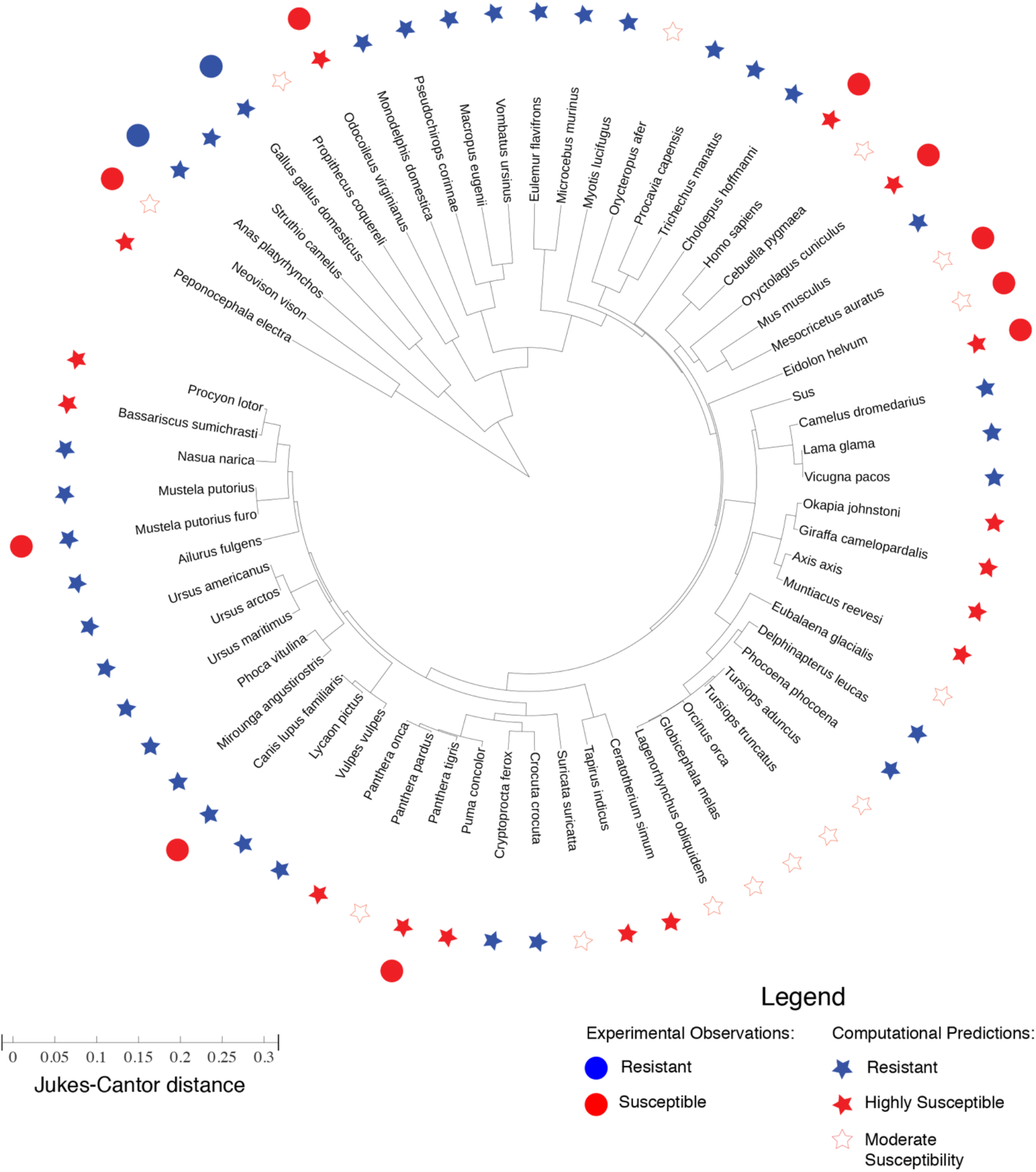
Phylogenic tree representing the evolutionary distance between ACE2 proteins of different species. The lengths in the radial direction denote the Jukes-Cantor distance (See Materials & Methods) as a measure of evolutionary distance between any two ACE2 proteins. The experimental observation of SARS-CoV-2 resistance/susceptibility are plotted alongside the computational predictions for resistance/susceptibility based on our frustration analysis of the ACE2-spike complex. See the Legend for more details. There is a consistency between the computational predictions and the experimental observations for mouse (*Mus musculus*), chicken (*Gallus gallus domesticus*)(Shi et al., 2020), duck (*Anas platyrhynchos*)(Shi et al., 2020), mink (*Neovison vison*)(Oude Munnink et al., 2021), bat (*Eidolon helvum*)(Kumakamba et al., 2020), Syrian golden hamster (*Mesocricetus auratus*)(Sia et al., 2020), tiger (*Panthera tigris*), white-tailed deer (*Odocoileus virginianus*)(Palmer et al., 2021), European rabbit (*Oryctolagus cuniculus*)(Mykytyn et al., 2020), and pig (*Sus scrofa*)(BS et al., 2021). However, apparent inconsistencies are found for ferret (*Mustela putorius furo*)(Shi et al., 2020) and dog (*Canis lupis familiaris*) (Shi et al., 2020; Sit et al., 2020)—however, SARS-Cov-2 has only been observed to replicate in the upper respiratory tract of ferrets(Shi et al., 2020), and viral replication has been observed to be low in dogs(Shi et al., 2020).

We further apply this analysis for identifying frustrated outliers in our other modeled complexes, thereby predicting whether a particular animal is susceptible to COVID-19. A detailed summary of our results is shown in Figure 2, which includes experimental observations that corroborate or are inconsistent with our predictions. Other animals that have been experimentally observed to be susceptible to COVID-19, such as mink (*Neovison vison*) (Oude Munnink et al., 2021), and Syrian golden hamster (*Mesocricetus auratus*)(Sia et al., 2020), are identified as being moderately susceptible by our computational approach. Coronavirus consensus PCR-primer sequences have been detected with high frequency in populations of straw-colored fruit bats (*Eidolon helvum*)(Kumakamba et al., 2020), which have been predicted to exhibit moderate susceptibility by the computational approach.

Taken together, our findings summarized in Figure 2 show that an energy landscape-based approach can identify the molecular origins of COVID-19 susceptibility and resistance. Experimental observations corroborate 10 out of 12 of the computational predictions. One apparent inconsistency between the predictions and experimental observation regards the susceptibility of ferrets (*Mustela putorius furo*) which have been observed to replicate SARS-Cov-2 specifically in their upper respiratory tract(Shi et al., 2020). Our analysis of the ferret ACE2-spike complex reveals a single highly frustrated inter-protein interaction between 34Y of ACE2 and 403R of the spike protein. However, it has been noted(Damas et al., 2020; Shi et al., 2020) that ferrets have a unique respiratory biology, which may offer an explanation for this apparent discrepancy. Another apparent inconsistency is observed with our predictions for dogs (*Canis lupus familiaris*). Our frustration analysis predicts two highly frustrated inter-protein contacts within the ACE2/spike complex: 33Y of ACE2 with 417K of the spike and 325E of the ACE2 with 502G of the spike. Yet the susceptibility of dogs still remains somewhat controversial—while viral susceptibility and the production of antibody responses have been detected in dogs(Sit et al., 2020), viral replication has been reported to be poor(Shi et al., 2020).

Our energy landscape-based predictions for COVID-19 susceptibility are closely related to similar approaches that examined sequence differences in ACE2 sequences of different animals in the context of a structural model of the ACE2/spike complex(Damas et al., 2020; Lam et al., 2020; Luan et al., 2020). Frustration analysis yields a benefit to computational estimates of binding affinity because it compares the interaction energies between ACE2 and the spike glycoprotein with respect to alternative configurations (i.e., decoys) to assess how favorable a particular interaction is in the binding interface. In particular, the majority of our predictions are consistent with those of Damas et al(Damas et al., 2020), which makes predictions that are consistent with the same 10 out of 12 experiment observations that are highlighted in Figure 2. However, validation of the different models that exists is limited by the relatively small number of confirmed cases of COVID-19 in animals.

By in large, we find that our simple model appears to be consistent with many experimental observations of COVID-19 infections across different animals despite only considering the ACE2-spike protein interaction.

## Conclusion

The COVID-19 pandemic and the spread of other coronaviruses in recent years requires an indirect approach to understanding the molecular determinants behind susceptibility and resistance. Here, we constructed structural models of the ACE2-spike glycoprotein complex for a wide range of animals with ACE2 receptors. Using an energy landscape theory-based analysis we are able to uncover specific inter-protein interactions between the ACE2 and spike that appear to confer COVID-19 resistance. Our predictions appear to be consistent with many of the experimental observations regarding animal susceptibility, providing a structural explanation to those observations.

Our analysis reveals that the evolutionary distance between ACE2 proteins is not sufficient to predict COVID-19 susceptibility (Figure 2). Rather, an energy landscape-based analysis appears necessary to assess the interactions between the ACE2 protein and the SARS-Cov-2 spike glycoprotein.

## Acknowledgements

The authors would like to thank Matthew MacManes, Erez Aiden, and Olga Dudchenko for helpful discussions and providing data and resources through the DNA Zoo. This work was supported by the Center for Theoretical Biological Physics sponsored by the National Science Foundation NSF Grant PHY-2019745. J.N.O. was also supported by the NSF-CHE-1614101 and by the Welch Foundation (Grant C-1792). J.N.O. is a Cancer Prevention and Research Institute of Texas. Scholar in Cancer Research.

---

**Figure S1.**
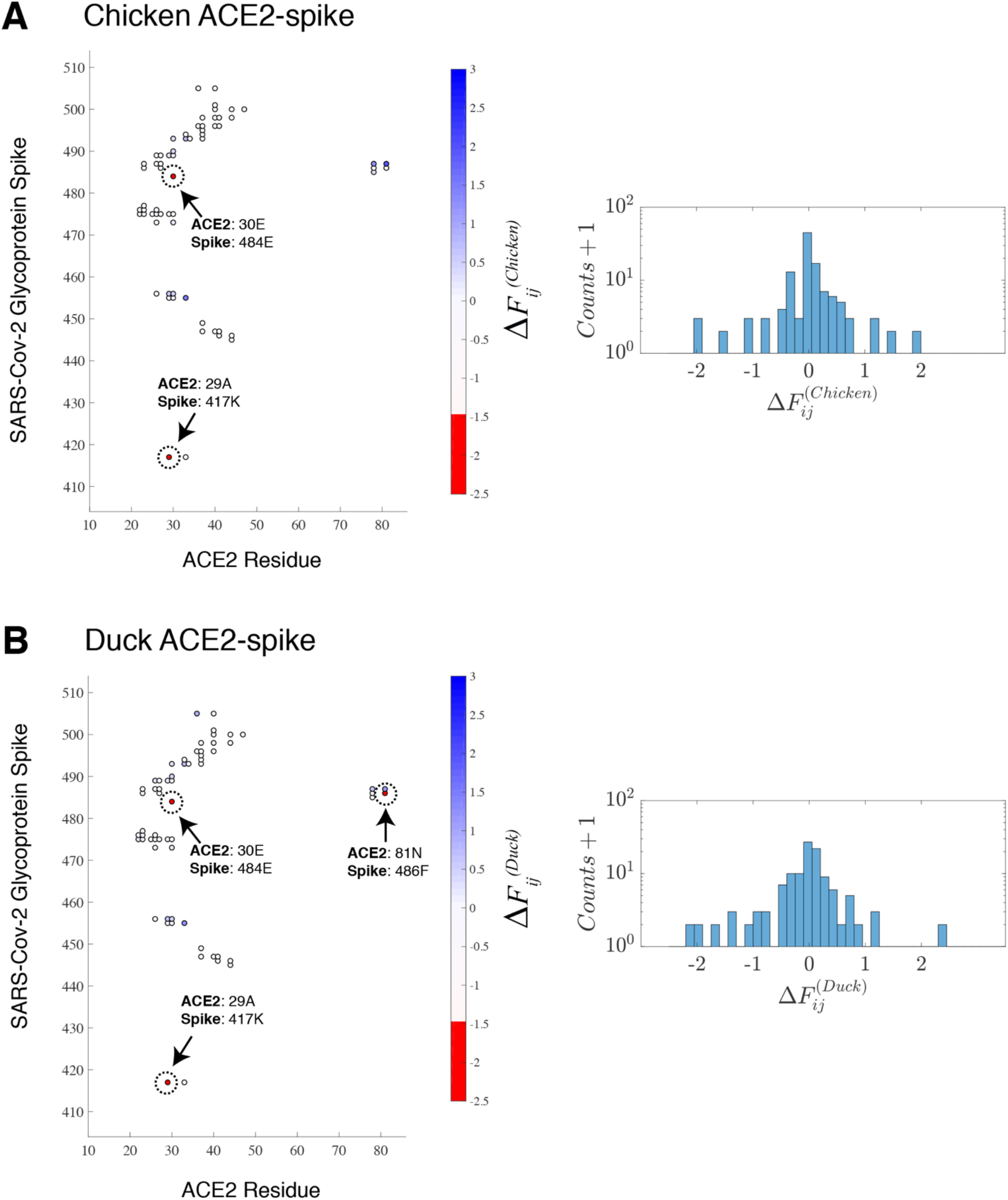
Comparative analysis of the frustration indices with respect to those observed in human ACE2-spike complex for additional examples of animals with known COVID-19 resistance. The configurational frustration index relative to the frustration in the human ACE2-spike complex is shown for (A) chicken (*Gallus gallus domesticus*) and (B) duck (*Anas platyrhynchos*) on a contact map illustrating select contacts between the SARS-Cov-2 spike and the ACE2 protein. Corresponding histograms of the frustration index between all contacts between the ACE2 and spike protein are also shown. Animals that resist COVID-19 appear to have frustrated outliers that represent highly unfavorable residue interactions compared to the human ACE2-spike complex (i.e., 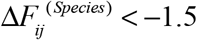). For the chicken, two highly frustrated interactions are identified: between (1) 30E of the ACE2 and 484E of the spike and (2) 29A of the ACE2 and 417K of the spike. For the duck, three highly frustrated interactions are identified: between (1) 30E of the ACE2 and 484E of the spike, (2) 29A of the ACE2 and 417K of the spike, and (3) 81N of the ACE2 and 486F of the spike. These frustrated interactions appear to confer COVID-19 resistance.

**Figure S2.**
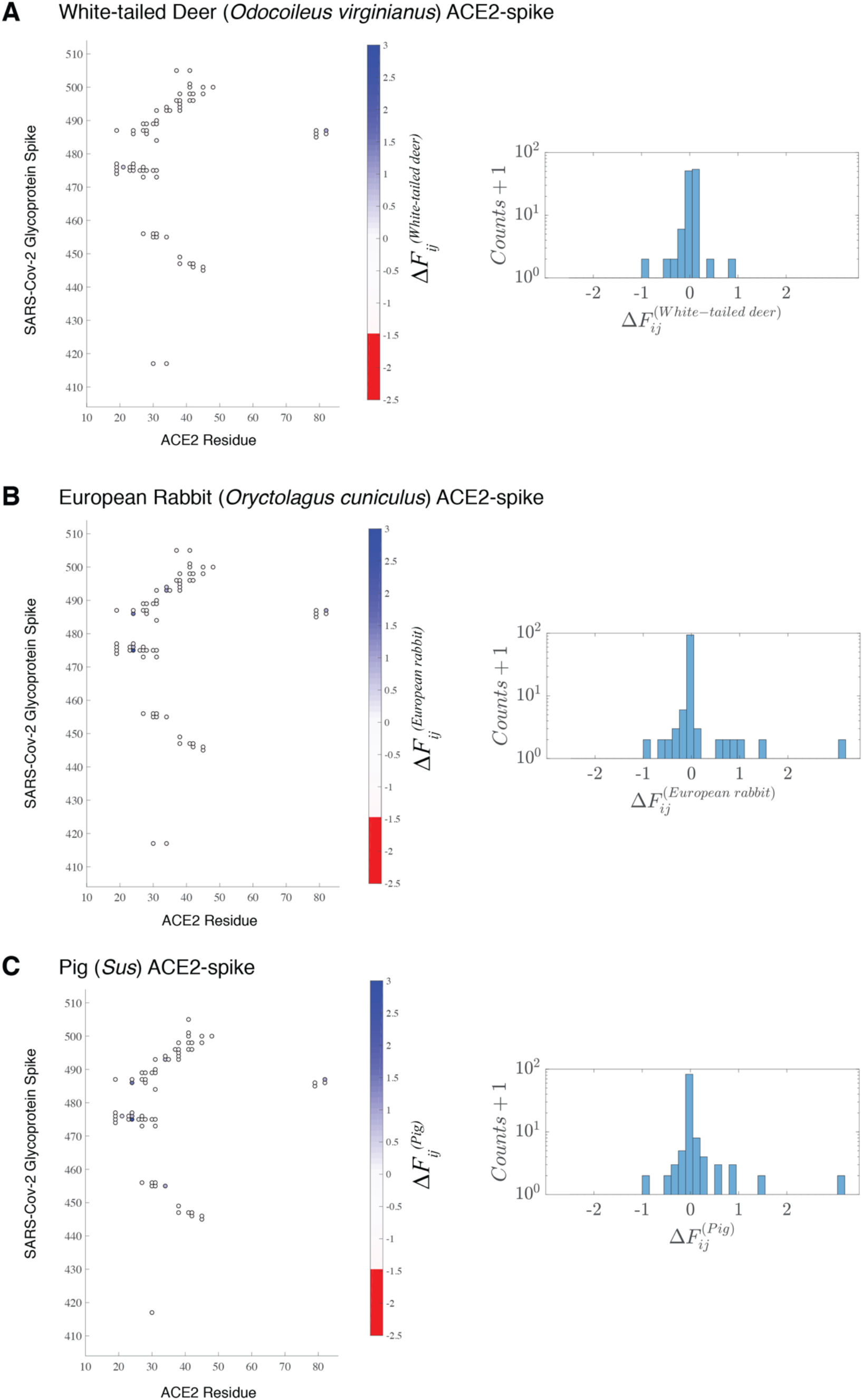
Comparative analysis of the frustration indices with respect to those observed in human ACE2-spike complex for additional examples of animals with known COVID-19 susceptibility. The configurational frustration index relative to the frustration in the human ACE2-spike complex is shown for (A) white-tailed deer (*Odocoileus virginianus*), (B) European rabbit (*Oryctolagus cuniculus*), and (C) pig (*Sus*) on a contact map illustrating select contacts between the SARS-Cov-2 spike and the ACE2 protein. Corresponding histograms of the frustration index between all contacts between the ACE2 and spike protein are also shown. Animals that are susceptible to COVID-19 appear to have comparable levels of frustration compared to human ACE2-spike complex.

## Notes

### Competing Interest Statement

The authors have declared no competing interest.

### Summary of Updates

SI Figures added

## References

Becker, D. J., Albery, G. F., Sjodin, A. R., Poisot, T., Dallas, T. A., Eskew, E. A., … Carlson, C. J. (2020). Predicting wildlife hosts of betacoronaviruses for SARS-CoV-2 sampling prioritization: a modeling study. bioRxiv, 2020.2005.2022.111344. doi:10.1101/2020.05.22.111344

BS, P., G, S., MM, P., C, E.-H., E, M., P, M., & CE, L. (2021). Susceptibility of Domestic Swine to Experimental Infection with Severe Acute Respiratory Syndrome Coronavirus 2. Emerg Infect Dis., 27(1), 104–112. doi:https://dx.doi.org/10.3201/eid2701.203399

Damas, J., Hughes, G. M., Keough, K. C., Painter, C. A., Persky, N. S., Corbo, M., … Lewin, H. A. (2020). Broad host range of SARS-CoV-2 predicted by comparative and structural analysis of ACE2 in vertebrates. Proceedings of the National Academy of Sciences, 117(36), 22311. doi:10.1073/pnas.2010146117

Davtyan, A., Schafer, N. P., Zheng, W., Clementi, C., Wolynes, P. G., & Papoian, G. A. (2012). AWSEM-MD: Protein Structure Prediction Using Coarse-Grained Physical Potentials and Bioinformatically Based Local Structure Biasing. The Journal of Physical Chemistry B, 116(29), 8494–8503. doi:10.1021/jp212541y

Dudchenko, O., Batra, S. S., Omer, A. D., Nyquist, S. K., Hoeger, M., Durand, N. C., … Aiden, E. L. (2017). De novo assembly of the <em>Aedes aegypti</em> genome using Hi-C yields chromosome-length scaffolds. Science, 356(6333), 92. doi:10.1126/science.aal3327

Ferreiro, D. U., Hegler, J. A., Komives, E. A., & Wolynes, P. G. (2007). Localizing frustration in native proteins and protein assemblies. Proceedings of the National Academy of Sciences, 104(50), 19819. doi:10.1073/pnas.0709915104

Frank, H. K., Enard, D., & Boyd, S. D. (2020). Exceptional diversity and selection pressure on SARS-CoV and SARS-CoV-2 host receptor in bats compared to other mammals. bioRxiv, 2020.2004.2020.051656. doi:10.1101/2020.04.20.051656

Gautam, A., Kaphle, K., Shrestha, B., & Phuyal, S. (2020). Susceptibility to SARS, MERS, and COVID-19 from animal health perspective. Open Vet J, 10(2), 164–177. doi:10.4314/ovj.v10i2.6

Goldstein, J. (2020). Bronx Zoo Tiger Is Sick With the Coronavirus. New York Times. Retrieved from https://www.nytimes.com/2020/04/06/nyregion/bronx-zoo-tiger-coronavirus.html

Kumakamba, C., Niama, F. R., Muyembe, F., Mombouli, J.-V., Kingebeni, P. M., Nina, R. A., … Lange, C. E. (2020). Coronavirus surveillance in Congo basin wildlife detects RNA of multiple species circulating in bats and rodents. bioRxiv, 2020.2007.2020.211664. doi:10.1101/2020.07.20.211664

Lam, S. D., Bordin, N., Waman, V. P., Scholes, H. M., Ashford, P., Sen, N., … Orengo, C. A. (2020). SARS-CoV-2 spike protein predicted to form complexes with host receptor protein orthologues from a broad range of mammals. Scientific Reports, 10(1), 16471. doi:10.1038/s41598-020-71936-5

Li, W., Moore, M. J., Vasilieva, N., Sui, J., Wong, S. K., Berne, M. A., … Farzan, M. (2003). Angiotensin-converting enzyme 2 is a functional receptor for the SARS coronavirus. Nature, 426(6965), 450–454. doi:10.1038/nature02145

Luan, J., Lu, Y., Jin, X., & Zhang, L. (2020). Spike protein recognition of mammalian ACE2 predicts the host range and an optimized ACE2 for SARS-CoV-2 infection. Biochemical and Biophysical Research Communications, 526(1), 165–169. doi:https://doi.org/10.1016/j.bbrc.2020.03.047

Madeira, F., Park, Y. M., Lee, J., Buso, N., Gur, T., Madhusoodanan, N., … Lopez, R. (2019). The EMBL-EBI search and sequence analysis tools APIs in 2019. Nucleic Acids Research, 47(W1), W636–W641. doi:10.1093/nar/gkz268

Martínez-Hernández, F., Isaak-Delgado, A. B., Alfonso-Toledo, J. A., Muñoz-García, C. I., Villalobos, G., Aréchiga-Ceballos, N., & Rendón-Franco, E. (2020). Assessing the SARS-CoV-2 threat to wildlife: Potential risk to a broad range of mammals. Perspectives in Ecology and Conservation. doi:https://doi.org/10.1016/j.pecon.2020.09.008

Melin, A. D., Janiak, M. C., Marrone, F., Arora, P. S., & Higham, J. P. (2020). Comparative ACE2 variation and primate COVID-19 risk. Communications Biology, 3(1), 641. doi:10.1038/s42003-020-01370-w

Muñoz-Fontela, C., Dowling, W. E., Funnell, S. G. P., Gsell, P.-S., Riveros-Balta, A. X., Albrecht, R. A., … Barouch, D. H. (2020). Animal models for COVID-19. Nature, 586(7830), 509–515. doi:10.1038/s41586-020-2787-6

Mykytyn, A. Z., Lamers, M. M., Okba, N. M. A., Breugem, T. I., Schipper, D., van den Doel, P. B., … Haagmans, B. L. (2020). Susceptibility of rabbits to SARS-CoV-2. bioRxiv, 2020.2008.2027.263988. doi:10.1101/2020.08.27.263988

Onuchic, J. N., Luthey-Schulten, Z., & Wolynes, P. G. (1997). THEORY OF PROTEIN FOLDING: The Energy Landscape Perspective. Annual Review of Physical Chemistry, 48(1), 545–600. doi:10.1146/annurev.physchem.48.1.545

Onuchic, J. N., & Wolynes, P. G. (2004). Theory of protein folding. Current Opinion in Structural Biology, 14(1), 70–75. doi:https://doi.org/10.1016/j.sbi.2004.01.009

Oude Munnink, B. B., Sikkema, R. S., Nieuwenhuijse, D. F., Molenaar, R. J., Munger, E., Molenkamp, R., … Koopmans, M. P. G. (2021). Transmission of SARS-CoV-2 on mink farms between humans and mink and back to humans. Science, 371(6525), 172. doi:10.1126/science.abe5901

Palmer, M. V., Martins, M., Falkenberg, S., Buckley, A., Caserta, L. C., Mitchell, P. K., … Diel, D. G. (2021). Susceptibility of white-tailed deer (<em>Odocoileus virginianus</em>) to SARS-CoV-2. bioRxiv, 2021.2001.2013.426628. doi:10.1101/2021.01.13.426628

Parra, R. G., Schafer, N. P., Radusky, L. G., Tsai, M.-Y., Guzovsky, A. B., Wolynes, P. G., & Ferreiro, D. U. (2016). Protein Frustratometer 2: a tool to localize energetic frustration in protein molecules, now with electrostatics. Nucleic Acids Research, 44(W1), W356–W360. doi:10.1093/nar/gkw304

Shang, J., Ye, G., Shi, K., Wan, Y., Luo, C., Aihara, H., … Li, F. (2020). Structural basis of receptor recognition by SARS-CoV-2. Nature, 581(7807), 221–224. doi:10.1038/s41586-020-2179-y

Shi, J., Wen, Z., Zhong, G., Yang, H., Wang, C., Huang, B., … Bu, Z. (2020). Susceptibility of ferrets, cats, dogs, and other domesticated animals to SARS–coronavirus 2. Science, 368(6494), 1016. doi:10.1126/science.abb7015

Sia, S. F., Yan, L.-M., Chin, A. W. H., Fung, K., Choy, K.-T., Wong, A. Y. L., … Yen, H.-L. (2020). Pathogenesis and transmission of SARS-CoV-2 in golden hamsters. Nature, 583(7818), 834–838. doi:10.1038/s41586-020-2342-5

Sit, T. H. C., Brackman, C. J., Ip, S. M., Tam, K. W. S., Law, P. Y. T., To, E. M. W., … Peiris, M. (2020). Infection of dogs with SARS-CoV-2. Nature, 586(7831), 776–778. doi:10.1038/s41586-020-2334-5

The UniProt, C. (2021). UniProt: the universal protein knowledgebase in 2021. Nucleic Acids Research, 49(D1), D480–D489. doi:10.1093/nar/gkaa1100

Wang, Q., Zhang, Y., Wu, L., Niu, S., Song, C., Zhang, Z., … Qi, J. (2020). Structural and Functional Basis of SARS-CoV-2 Entry by Using Human ACE2. Cell, 181(4), 894–904.e899. doi:https://doi.org/10.1016/j.cell.2020.03.045

Waterhouse, A., Bertoni, M., Bienert, S., Studer, G., Tauriello, G., Gumienny, R., … Schwede, T. (2018). SWISS-MODEL: homology modelling of protein structures and complexes. Nucleic Acids Research, 46(W1), W296–W303. doi:10.1093/nar/gky427

Yan, R., Zhang, Y., Li, Y., Xia, L., Guo, Y., & Zhou, Q. (2020). Structural basis for the recognition of SARS-CoV-2 by full-length human ACE2. Science, 367(6485), 1444. doi:10.1126/science.abb2762

Zhou, P., Yang, X.-L., Wang, X.-G., Hu, B., Zhang, L., Zhang, W., … Shi, Z.-L. (2020). A pneumonia outbreak associated with a new coronavirus of probable bat origin. Nature, 579(7798), 270–273. doi:10.1038/s41586-020-2012-7

